# Intergenic RNA mainly derives from nascent transcripts of known genes

**DOI:** 10.1101/2020.01.08.898478

**Authors:** Agostini Federico, Zagalak Julian, Attig Jan, Ule Jernej, Nicholas M. Luscombe

**Author notes:** Senior authors (Ule Jernej, Nicholas M. Luscombe).

## Abstract

**Background:** Eukaryotic genomes undergo pervasive transcription, leading to the production of many types of stable and unstable RNAs. Transcription is not restricted to regions with annotated gene features but includes almost any genomic context. Currently, the source and function of most RNAs originating from intergenic regions in the human genome remains unclear.

**Results:** We hypothesised that many intergenic RNA can be ascribed to the presence of as-yet unannotated genes or the ‘fuzzy’ transcription of known genes that extends beyond the annotated boundaries. To elucidate the contributions of these two sources, we assembled a dataset of >2.5 billion publicly available RNA-seq reads across 5 human cell lines and multiple cellular compartments to annotate transcriptional units in the human genome. About 80% of transcripts from unannotated intergenic regions can be attributed to the fuzzy transcription of existing genes; the remaining transcripts originate mainly from putative long non-coding RNA loci that are rarely spliced. We validated the transcriptional activity of these intergenic RNA using independent measurements, including transcriptional start sites, chromatin signatures, and genomic occupancies of RNA polymerase II in various phosphorylation states. We also analysed the nuclear localisation and sensitivities of intergenic transcripts to nucleases to illustrate that they tend to be rapidly degraded either ‘on-chromatin’ by XRN2 or ‘off-chromatin’ by the exosome.

**Conclusions:** We provide a curated atlas of intergenic RNAs that distinguishes between alternative processing of well annotated genes from independent transcriptional units based on the combined analysis of chromatin signatures, nuclear RNA localisation and degradation pathways.

## Background

Studies estimate that up to 85% of the human genome is pervasively transcribed by RNA polymerase II (Pol II), resulting in a plethora of RNA products [1–4]. Many of these transcripts belong to well established categories, such as messenger RNAs (mRNAs) which are characterised by the presence of 5’ cap, coding sequence (CDS) and poly(A) tail. Other transcripts are categorised as long non-coding RNAs (lncRNAs), generally defined as RNA molecules longer than 200 nt with little coding potential. Currently, lncRNAs are divided into three major groups depending on their genomic location relative to protein-coding genes: promoter upstream transcripts (PROMPTs), produced up to 2.5 kb upstream of active transcription start sites (TSSs) [5]; enhancer RNAs (eRNAs), bi-directionally transcribed from enhancer DNA elements [6,7]; and large intervening non-coding RNAs (lincRNAs), located in intergenic regions, distal from protein-coding genes and regulated as independent transcriptional units [8]. Gene and transcript annotations for the human genome are continuously updated and their assignment to specific biotype categories can change across reference databases [9]. In particular, in the past decade, efforts towards the identification and characterisation of novel lncRNA genes have been made, either through computational predictions or functional assays [10,11]. Despite such endeavours however, a marked proportion of RNA-seq reads from human cells still map to unannotated, ostensibly intergenic portions of the human genome. It is therefore often challenging to understand whether such reads originate from independent transcription units or are associated with annotated genes.

Many well-characterised lncRNAs, such as the X-inactive specific transcript *Xist* [12], share processing features (*e.g.*, 5’ m^7^G cap and poly(A) tail) with mRNAs [8] and have specific, experimentally validated functions. However, the majority of lncRNA gene loci might not function through their resulting products, but rather through the act of transcription itself, which for instance can affect the expression of neighbouring genes [13–15]. In support of this view, studies have highlighted how ncRNA genes are associated with early transcriptional termination of Pol II and their products undergo rapid post-transcriptional degradation [3,16–19], thus explaining their low nuclear abundance. Further, recent studies indicate a possible scenario in which nascent transcripts from protein-coding genes play a similar role by regulating chromatin remodelling [20]. For example, the binding of Polycomb repressive complex 2 (PRC2) to genomic targets was initially ascribed to a specific set of lncRNAs [21–24]. However, it was later shown that PRC2 also bind nascent, unspliced mRNAs, which sequester the complex, thus preventing gene silencing [25–28].

In addition to mRNAs and lncRNAs described above, downstream of gene transcripts (DoGs) arise when Pol II terminates far downstream of the ends of genes [29]. These readthrough transcripts appear to be linked to stress conditions, such as osmotic and oxidative stress [29,30]. It remains unclear whether transcription of DoGs has any gene regulatory function, but possible roles range from antisense-mediated gene expression control [31] to maintenance of local open chromatin structure. Moreover, their regulation remains largely unknown. Nevertheless, the existence of DoGs increases the complexity of transcriptome annotation, posing additional challenges to the understanding of function and regulation of intergenic transcripts.

In a recent study [32], we performed RNA-seq of the nuclear and cytoplasmic compartments of untreated HeLa cells and found that an unexpectedly large fraction (7.63%) of nuclear RNA-seq reads derived from intergenic genomic regions. Since the majority of these reads (60.3%) could not be detected in the cytoplasmic samples, here, we seek to investigate their transcriptional origin. We developed a computational method to identify and classify sources of intergenic transcription. We investigate their characteristics, expression patterns and epigenetic environment. Specifically, we observe that the largest fraction of intergenic RNA corresponds to DoGs, upstream of gene transcripts (UoGs), which likely result from alternative TSSs upstream of annotated genes, and linker of genes (LoGs), which are DoGs that continue into the neighbouring gene body. We find that most intergenic RNA is generated during transcription associated with annotated genes, and is confined to chromatin due to efficient degradation of DoGs and LoGs by XRN2, and UoGs by the exosome. Most remaining intergenic RNA corresponds to poorly spliced lncRNAs that are degraded by the exosome. We conclude that most of the unannotated intergenic RNAs are the consequence of non-productive transcription associated with known genes, which are rapidly removed through cellular quality control mechanisms.

## Results

### Identification of intergenic transcriptional units

To gain a comprehensive overview of the transcriptional landscape, we identified 38 publicly available datasets containing chromatin and nuclear fractionated RNA-seq samples. These cover 5 human cell lines (HeLa, HEK293, HepG2, K562, HCT116) and four subcellular fractions (cytosolic, nuclear, chromatin and nucleoplasm). Initial processing and mapping to the human genome yielded >2.5 billion uniquely mapped reads (Figure 1, Supplementary Table 1). We employed StringTie [33] to generate preliminary annotations of the transcriptional units expressed within each dataset. We then merged the results into a comprehensive transcriptomic assembly across the entire dataset and also included all genes present in the GENCODE reference annotation [34]. Finally, we employed a custom pipeline (Materials & Methods) to annotate transcripts expressed in intergenic regions and to define their relationship with annotated genes (Figure 1 and 2A; see Materials and Methods). We defined transcriptional units (TU) as products of transcription from intergenic portions of the genome, which can either take place as an independent event or in association with features in the reference annotation.

**Figure 1.**
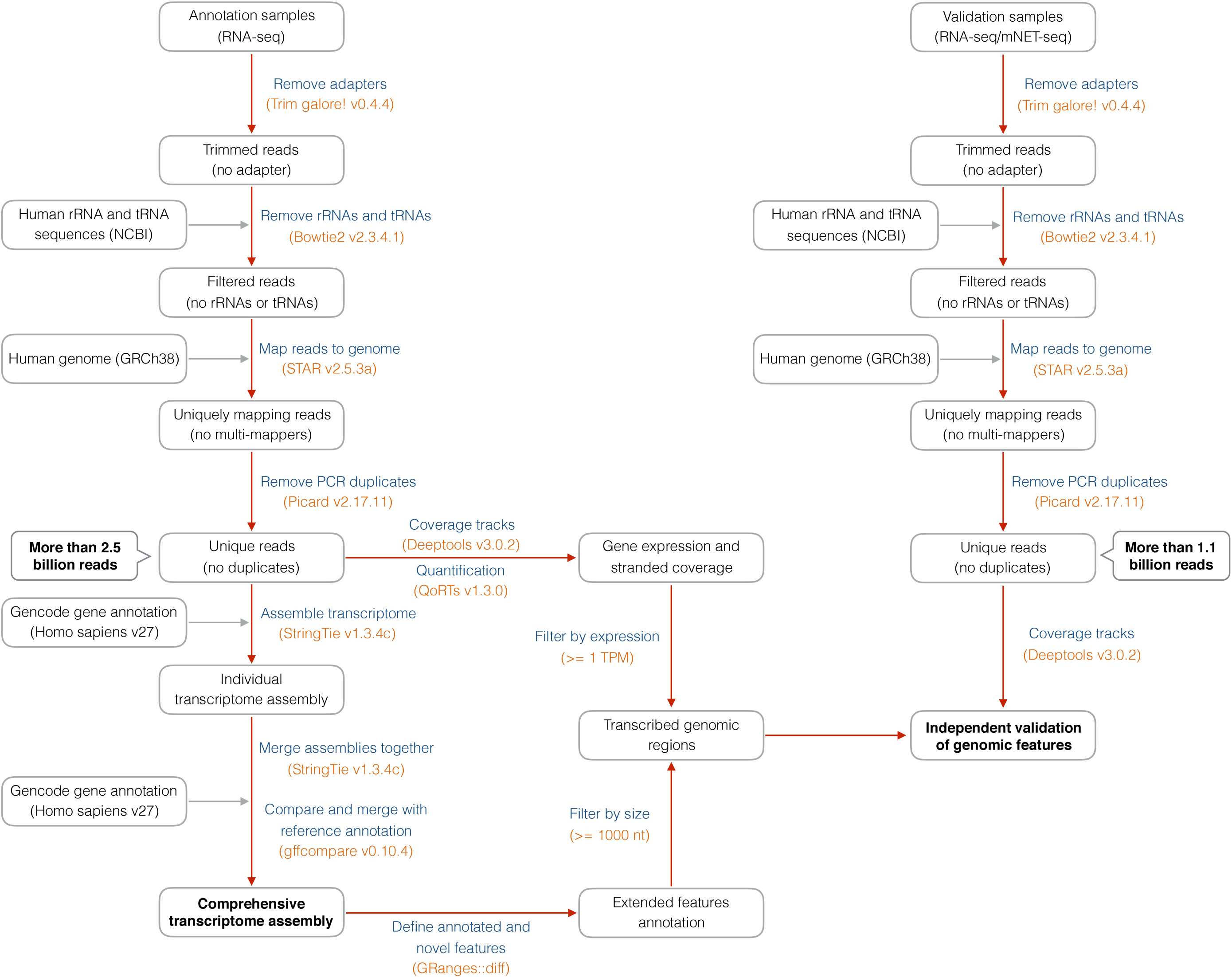
Flow chart of data analysis pipeline. Schematic describing the main data processing steps, intermediate and final outputs of the analysis pipeline, applied to RNA-seq (left side) and other sequencing (NET-seq, right side) data. Procedures (blue) and tools (orange) are indicated.

**Figure 2.**
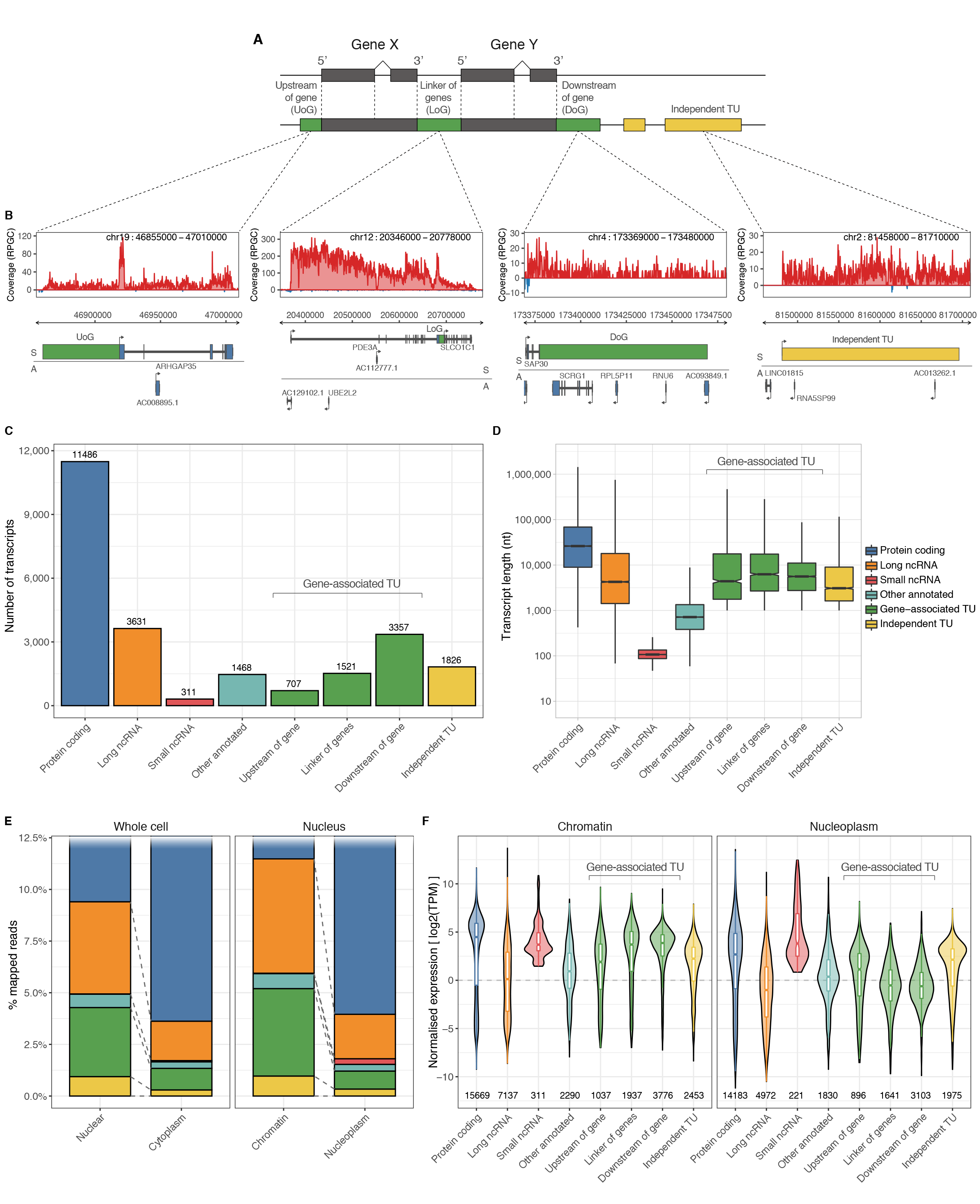
General features of newly identified transcriptional units (TUs). A) Schematic representation of the gene-associated (green) and independent (yellow) transcriptional units annotated in this study. B) Upper panels, genome-browser views of nuclear RNA-seq signals in HeLa cells for example TUs (red and blue indicate RNA-seq reads mapping to the sense and antisense strands respectively) Lower panels, genomic annotations of pre-existing genes and newly identified TUs; horizontal line divides features on the sense (S) and antisense (A) orientations. Coverage is reported at 1x depth (reads per genome coverage, RPGC). C) Comparison of number of annotated and newly identified transcripts detected in the current RNA-seq dataset (TPM >= 1). D) Comparison of transcript lengths. E) Proportions of uniquely mapping RNA-seq reads originating from different transcript types for whole cell (left) and nuclear (right) subcellular fractions of HeLa cells. F) Distributions of expression levels of annotated and newly identified TUs for the chromatin-associated (left panel) and nucleoplasm (right panel) subcellular fractions of HeLa cells.

We classified TUs into two broad groups based on their genomic location relative to existing gene annotations (Figure 2A,B). (i) Gene-associated TUs are those showing continuity of transcription from the body of annotated genes. These were further divided into upstream of gene (UoG), downstream of gene (DoG) and linker of genes TUs (LoG). (ii) Independent TUs, which are >10 kb away from existing gene annotations and so classed as purely intergenic. Though we use the term ‘independent TUs’ for the sake of clarity, it is possible that some might in the future end up annotated as new genes that produce functional non-coding RNAs or perhaps even protein-coding mRNAs. In total, we classified 7,411 TUs covering ~5.6% of the human genome (Figure 2C). Both gene-associated and independent TUs are of comparable lengths to previously annotated long non-coding RNA genes, ranging from 1kb (the minimum length threshold for a TU) to hundreds of kb (Figure 2D).

Assessing expression levels in HeLa cells, it is apparent that the relative abundance of TUs is higher in the nucleus (4.28% of mapped reads) compared with the cytoplasm (1.34% of mapped reads; Figure 2E). Moreover within the nucleus, TUs tend to be chromatin-associated (5.2% of mapped reads) rather than the nucleoplasm (1.21%) (Figure 2E). Intriguingly, we noticed that about 80% of these reads mapped to features linked to transcription of previously annotated loci (*i.e.*, gene-associated TUs), while the remainder belong to independent TUs (Figure 2E). We also compared the normalised expression levels of TUs with annotated genes (Figure 2F). Protein-coding transcripts tend to be most highly expressed within the chromatin-associated and nucleoplasmic compartments; however, in these subcellular fractions, both gene-associated and independent TUs are more highly expressed than annotated lncRNAs. Additionally, we found independent TUs tend to undergo less splicing than lncRNAs (Figure S1B). In agreement with previous reports [29,35], DoGs and LoGs show the highest expression among TUs, suggesting that levels of transcription outside annotated loci primarily depend on the activity of annotated upstream features (Figure 2F).

To investigate the properties of TUs in greater detail, we focused further analysis to those with the strongest evidence:

- Both the gene-associated TU and neighbouring genes must have TPM expression ≥1 (Figure S1D) and length ≥5 kb (to avoid overlaps when assessing metaprofiles);
- UoG, LoG and DoG TUs must be associated with a protein-coding gene (Figure S1C), thus reducing the chance of including poorly annotated genes with relatively unreliable start and end genomic coordinates (e.g., pseudogenes);
- Independent TUs must be ≥10 kb from any annotated feature on the same strand orientation to ensure that they are not transcribed as part of a known gene (Figure S1A).

These filtering criteria left 1,604 gene-associated TUs (88 UoG, 1,329 DoG, 187 LoG) and 571 independent TUs. As controls, we paired gene-associated TUs with their corresponding protein-coding genes and we identified 3,462 lncRNA genes in a similar size range to independent TUs.

### Gene-associated transcription breaks gene boundaries

Next we sought to understand the transcriptional origins of gene-associated TUs. Here we focus on data from HeLa cells unless stated otherwise, as it is the cell type with the largest variety of measurements.

Figure 3 displays metaprofiles of diverse transcriptional measurements aligned to the start and ends of TUs and protein-coding genes (Fig. S2 for LoGs). All categories of TUs display clear RNA-seq coverage in the nuclear and chromatin-associated fractions, but in contrast to protein-coding genes, the signal is virtually lost in the cytoplasm (Fig. 3A and S2A). There is a clear jump in expression levels at the gene boundaries upon transition between the TU and associated gene, but TUs nonetheless display remarkably high relative expression levels in the nuclear and chromatin compartments.

**Figure 3.**
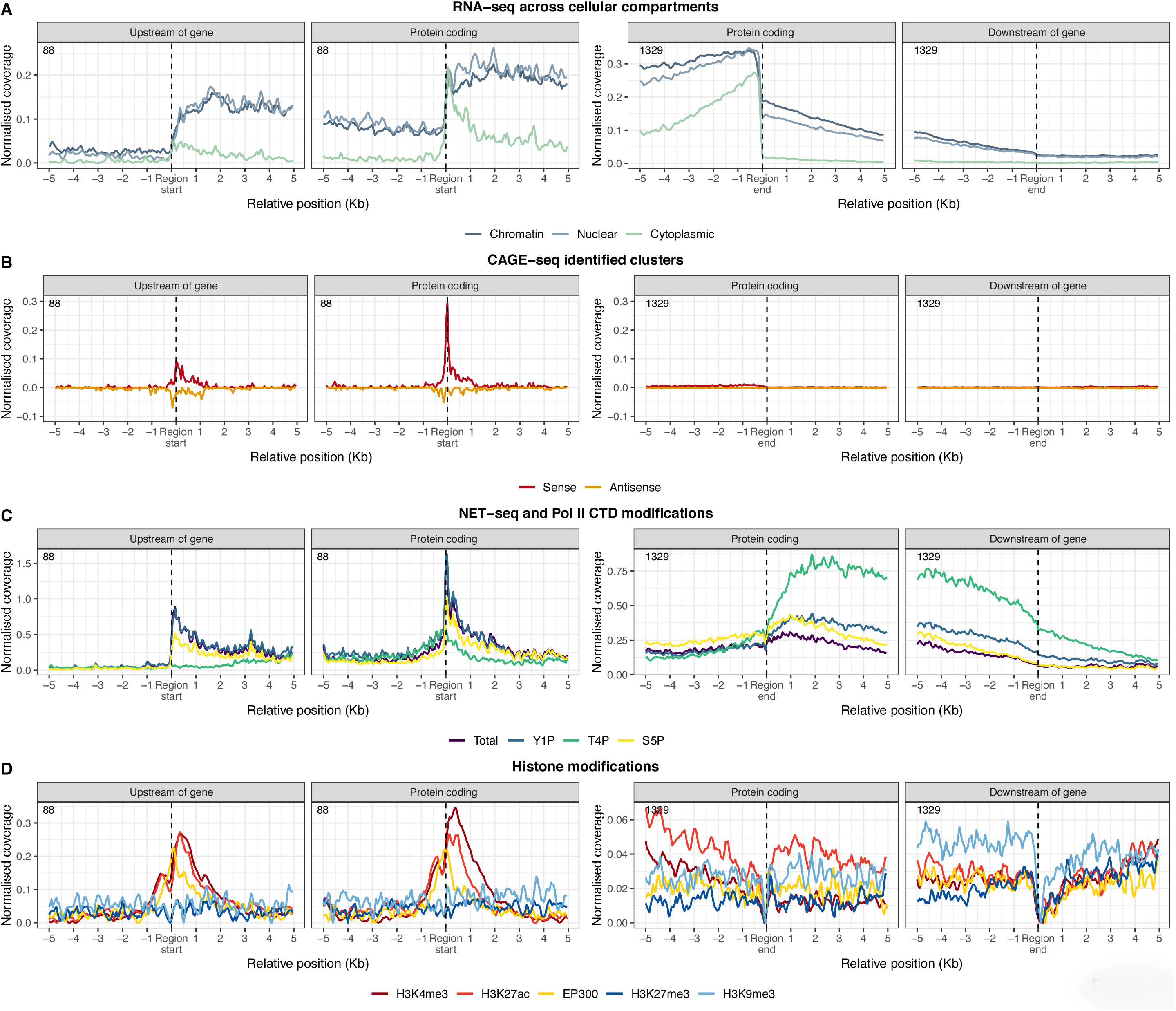
Meta-profiles of transcriptional measurements around gene-associated TUs. Meta-profiles of transcriptional measurements plotted relative to the start positions of UoGs and their associated protein-coding genes (left-hand panels), and relative to the end positions of DoGs and their associated genes (right-hand panels). A) RNA-seq measurements in different subcellular compartments; B) CAGE-seq measurements in the sense and antisense strands; C) NET-seq measurements for different Pol II CTD modifications; D) ChIP-seq measurements for histone marks and EP300 occupancies associated transcriptional activities.

Since the novel TUs are transcribed by RNA Pol II we asked if the unannotated TSSs initiating UoG transcription have previously been detected through cap analysis of gene expression (CAGE). To this end, we used annotated CAGE peaks derived from a large collection of cell lines and tissues [36,37]. UoGs display a slight enrichment of CAGE peaks at the start site, but we could hardly detect any signal for DoGs and LoGs (Figure 3B and S2B); this suggests that whereas UoGs show evidence of independent transcriptional initiation, DoGs and LoGs are most likely generated from transcriptional readthrough of the upstream gene. The modest CAGE signal for UoGs (detected for 48 out of 98 UoGs) suggest that they are not efficiently capped, in contrast to mRNAs initiating at annotated start sites of protein-coding genes (Figure 3B). This indicates that the majority of intergenic TUs might be designated as substrates for exonucleases and prone to degradation [38,39].

Figure 3C (and S3A) shows prominent Pol II occupancies at the start sites of UoGs, albeit at lower levels than at the TSS’ of associated genes. Together with the CAGE data, this possibly indicates the formation of a pre-initiation complex (PIC) and therefore the existence of unannotated, upstream TSSs. Active transcription of TUs is supported by mammalian native elongating transcript sequencing (NET-seq) data, which identifies nascent RNA fragments attached to transcriptionally engaged RNA Pol II [35]. NET-seq is capable of differentiating between distinct transcriptional stages by mapping nascent RNAs associated with different patterns of RNA Pol II C-terminal heptad repeat domain (CTD) phosphorylation. The annotated and UoG TSSs display similar NET-seq profiles, thus suggesting that the TUs are not the result of stochastic Pol II binding but rather the outcome of coordinated transcriptional initiation events. Indeed, the profile for tyrosine-1 (Y1P) phosphorylated Pol II - a hallmark of TSS-paused protein-coding gene transcripts [16] - displays the highest signal at the start positions of both UoGs and protein-coding genes, with the former having a less pronounced peak and a broader distribution of signal. Moreover, serine-5 (S5P) phosphorylated Pol II, which is mainly associated with TSS events such as co-transcriptional capping and early transcriptional elongation [40], follows a pattern similar to the total and Y1P profiles around these regions.

Threonine-4 (T4P) phosphorylation is a hallmark of terminating Pol II and causes a characteristic, NET-Seq signal near transcription end sites (TESs) of protein-coding genes [16]. Among protein-coding genes, the T4P profile peaks immediately after canonical TESs and remains high, while gradually decreasing towards the end of the associated DoG (Figure 3C). This observation implies that although Pol II is poised to terminate after encountering the canonical TES, actual Pol II detachment might occur several kilobases downstream. LoGs represent a special case, in which high T4P signal after the TES of the upstream gene is maintained throughout the intergenic space only to peak again at the TSS of the downstream gene (Figure S2C and S3B). This suggests either that transcription of LoGs joins two adjacent transcripts thereby generating a pseudo-bicistronic nascent RNAs or alternatively, that Pol II reaches the downstream gene and reinitiates transcription from a T4P state. In both cases, the downstream gene is potentially dependent on the transcription and by extension the promoter state of its upstream gene, thus implying the existence of co-regulation.

Finally, we examined the ChIP-seq profiles for four histone marks associated with transcriptional activity, as well as the histone acetyltransferase EP300 (Figure 3D and S4A). Epigenetic modifications such as H3K4me3 and H3K27ac, which are associated with active promoters and enhancers respectively [41,42], are enriched at both protein-coding and UoG start sites (Figure 3D). Furthermore, the tri-methylated forms of H3K27 and H3K9, commonly found at transcriptionally silenced regions [41,42], are depleted. Interestingly, the histone acetyltransferase EP300, which regulates transcription of genes via chromatin remodeling, shows a comparable enrichment in binding at both UoG TSSs and annotated TSSs (Figure 3D). EP300 is also known as a transcriptional coactivator, due to its ability to bind to transcription factors and the transcription machinery, and consequently activate transcription [41,42]. Therefore, the presence of this protein more than 5 kb (size used for selecting the intergenic features) upstream of the canonical TSS is intriguing, as it suggests that transcription from the upstream intergenic regions is not merely the consequence of stochastic initiation events, but rather a concerted and precisely regulated process.

### Transcription from deep intergenic regions

Next, we focused on independent TUs. We noticed a number of similarities between these elements and the 3,426 control lncRNAs. Specifically, for both classes we could detect RNA-seq signal upstream of the TSS and downstream of the TES (Figure 4A). This is probably due to the sub-optimal annotation of these reference positions, a challenging task considering the intrinsically low level of expression of such transcripts [43,44]. Interestingly, the CAGE signal displays equal enrichment in both orientations around the TSS of lncRNAs, possibly indicating that most of these RNAs originate from divergent transcription (Figure 4B). The NET-seq profiles show similar enrichment patterns for total RNA Pol II and the CTD modifications TSSs and TESs of lncRNAs and independent TUs (Figure 4C and S3C). Finally, the H3K9me3 and H3K27ac profiles around the TSSs of both lncRNAs and independent TUs resemble those of protein-coding genes and UoGs (Figure 4D and 3D), highlighting equivalent chromatin statuses. Thus based on the transcriptional and related measurements, independent TUs appear to be *bona fide* lincRNAs that eluded reference annotation.

**Figure 4.**
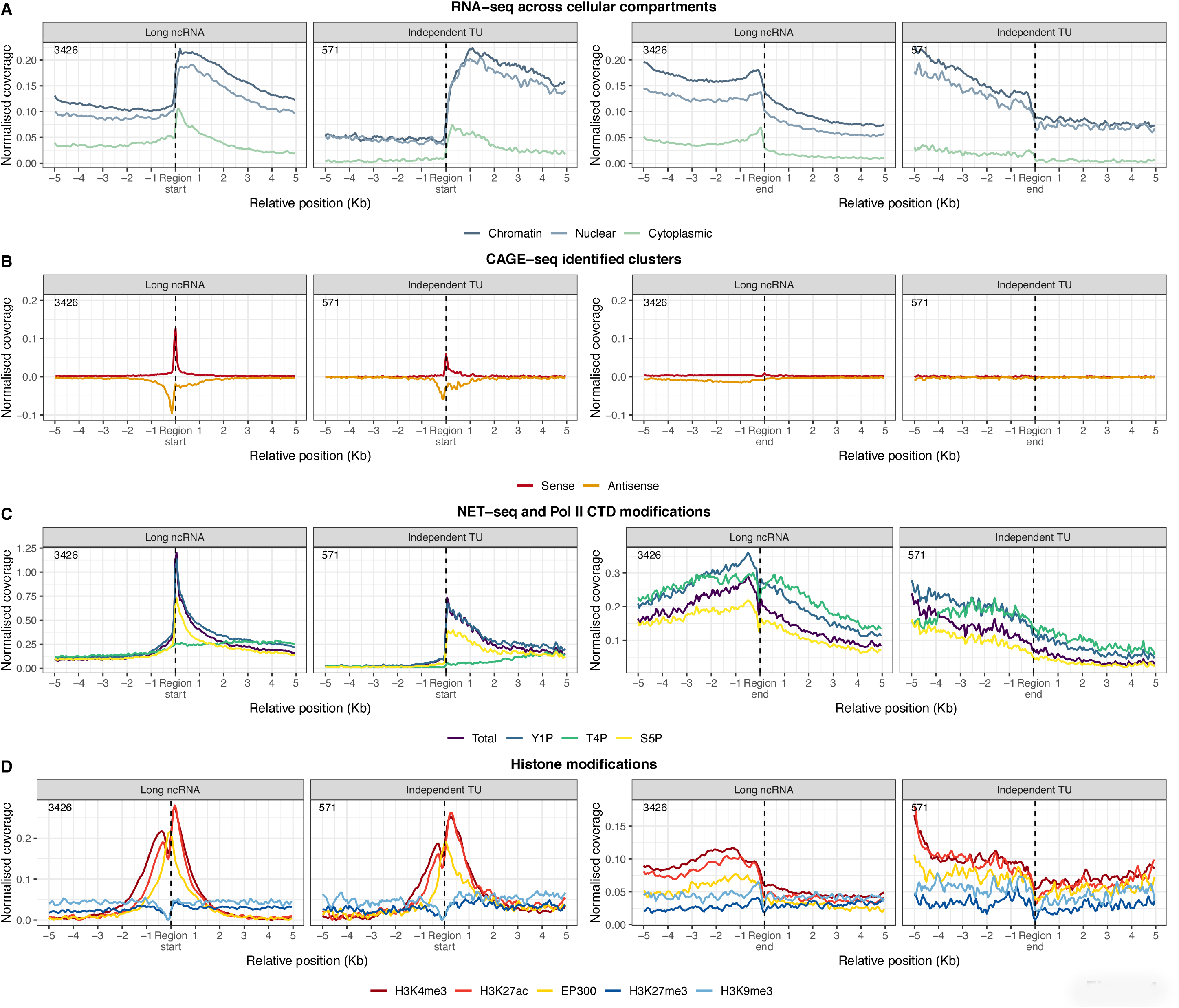
Meta-profiles of transcriptional measurements around independent TUs. Meta-profiles of transcriptional measurements plotted relative to the start and end positions of independent TUs and control long non-coding RNA genes. Panels A-D as in Figure 3.

### Rapid degradation of chromatin-associated intergenic RNAs

We showed that both gene-associated and independent TUs are widespread across the genome and their expression levels in the nucleus are comparable to those of annotated genes. Moreover, analysis of the transcribed loci did not highlight distinctive characteristics that explain why TUs are found only in the chromatin cellular compartment. Therefore, we hypothesised that there may be differences in the control of retention and stability of these transcripts.

First, we compared the expression levels of annotated RNAs and intergenic TUs between chromatin-associated and nucleoplasmic fractions. We found that unspliced protein-coding and long ncRNA transcripts tend to be equally distributed between the two fractions, whereas TUs, in particular DoGs and LoGs, are preferentially confined to the chromatin-associated fraction (Figure 5A and S5A).

**Figure 5.**
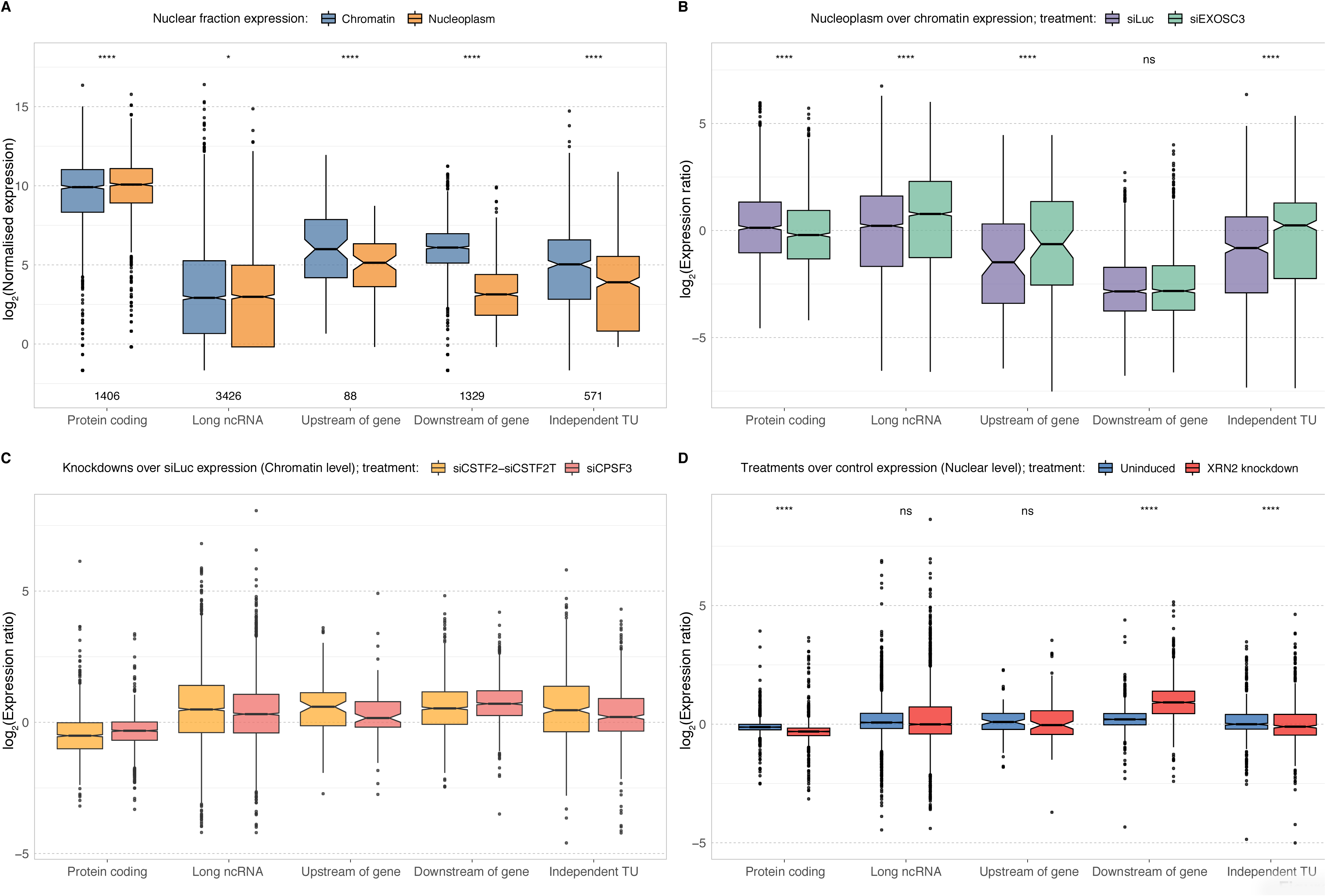
Impact of nuclease-depletion on TU expression. **A)** Expression levels of protein-coding genes and TUs in the chromatin and nucleoplasm fractions. B) Relative nucleoplasmic-to-chromatin expression levels in response to EXOSC3 knockdown and control siLuc treatments. C) Expression levels in CSTF2+CSTF2T and CPSF3 knockdowns relative to control in the chromatin fraction. D) Expression levels in XRN2 knockdown (via activation of auxin-inducible degron system) and basal (uninduced; minus auxin) treatments relative to unmodified XRN2 control in the nuclear fraction. P values were calculated using the two-sided Wilcoxon rank sum test, with asterisks indicating statistical significance at the following thresholds: ns (p > 0.05); * (p <= 0.05); ** (p <= 0.01); *** (p <= 0.001); **** (p <= 0.0001).

The scarcity of these transcripts in the nucleoplasm suggests that they exert their function, if any, bound to the chromatin fraction or that they are transcriptional byproducts that are rapidly degraded. We examined recently published RNA-seq data following knock-down or depletion of proteins involved in the processing and degradation of transcriptional products: specifically, EXOSC3 [16], CSTF2 (and its paralog CSTF2T), CPSF3 (also known as CPSF73) knockdowns in HeLa cells [35], and XRN2 depletion in HCT116 cells [45].

EXOSC3 is part of the RNA exosome complex; it possesses 3’ to 5’ exoribonuclease activity and it is involved in eliminating transcriptional byproducts. Known substrates include non-coding transcripts, such as promoter-upstream transcripts (PROMPTs), mRNAs with processing defects [46,47] and most prominently rRNA and snoRNAs, as part of their normal processing and maturation in the nucleolus [48]. The EXOSC3 knockdown had little or no effect on transcripts of protein-coding genes and their associated DoGs and LoGs; however, there is a marked effect on the stability of lncRNAs, UoGs and independent TUs in the nucleoplasmic fraction (Figure 5B and S5B). Moreover, the accumulation of these transcriptional products, caused by the loss of a functional nuclear RNA exosome, is more dramatic in the nucleoplasm than in the chromatin fraction, suggesting that they are generally targeted post-transcriptionally and cleared once they move away from the chromatin environment.

Since we observed a predominant chromatin retention (Figure 5A) and no effect of EXOSC3 knockdown on DoGs and LoGs (Figure 5B and S5B), we hypothesised that other mechanisms must regulate these TUs. We examined factors involved in processing the terminal regions of nascent transcripts: CSTF2 (and its paralog CSTF2T), implicated in 3’ end cleavage and polyadenylation of pre-mRNAs, CPSF3 (also known as CPSF73), a 3’ end-processing endonuclease, and XRN2, an exoribonuclease with 5’ to 3’ activity. Indeed, knockdowns of CPSF3 and of CSTF2+CSTF2T lead to increased levels of DoGs and LoGs (Figure 5C and S5C), suggesting that degradation of these transcripts is strongly dependent on the correct processing of the 3’ end of nascent transcripts. Downstream of cleavage at the polyA signal by the CPSF/CSTF complex, the remaining 3’ byproduct is depleted by the 3’ → 5’ exonuclease XRN2 [49]. Hence, we evaluated the expression of these transcripts in XRN2 depletion [45] to assess whether LoGs, like DoGs, are coupled with 3’ end processing of the upstream gene. XRN2 depletion greatly increased the expression of DoGs and LoGs, while leaving other transcript types unchanged (Figure 5D and S5D), thus indicating that XRN2 activity indeed regulates DoGs and LoGs abundance.

## Discussion

### Non-canonical transcription upstream of genes

To date, transcription upstream of canonical genes has been reported as a consequence of bidirectional transcription from neighbouring promoters or enhancers, with the transcript being generated in the antisense direction. In contrast to these transcripts, the UoGs identified here originate from the *same* strand as the associated downstream genes, thus limiting the possibility that these are products of enhancer- or promoter-derived divergent transcription. The presence of CAGE peaks on opposite strands around the beginning of these transcripts suggests that a minor fraction could instead originate from convergent transcription [50]. Either way, transcription close and across the canonical promoter region of the respective gene is expected to result in regulatory impact of UoG units, such as altering chromatin accessibility, or recruitment of Pol II and co-factors. Our study highlights examples in widespread used cell lines that can be studied in depth leveraging further genome-wide transcription data (such as HiC) or mechanistic analysis through genome editing.

### Non-canonical transcription downstream of genes

Studies have recently highlighted the presence of widespread transcription of intergenic regions downstream of protein-coding genes in mouse and human in response to heat shock, osmotic stress, or oxidative stress [29,30]. Although this form of transcriptional readthrough has been ascribed to the mammalian stress response, here we found evidence for such behaviour in unstimulated, normally proliferating cell lines. We observed two categories of readthrough, which are characterised by distinctive patterns of Pol II CTD phosphorylation. In the first group, DoGs arise from transcription of canonical genes that then continues for a few to hundreds of kilobases across intergenic space (DoG), as previously reported [16,29,35]. These are marked by the sharp increase in threonine 4 phosphorylation of Pol II (T4P) after the annotated TES, and the gradual and eventual loss of Pol II binding with distance. In the second group, Pol II continues transcribing to the next gene (LoG), thus hinting at the possibility of polycistronic transcription in higher eukaryotes [51]. In this case, the T4P signal does not fade, suggesting that most Pol II continues transcribing until it reaches the downstream gene. It is not clear whether Pol II proceeds uninterrupted through the next gene or reinitiates a separate transcriptional event. Co-regulation of genes in close proximity on the same chromosome has previously been described [52], and the existence of LoGs could be one of the factors explaining such observations.

### Functional consequences of non-canonical transcription on canonical genes

Although intergenic transcription has been commonly considered a consequence of pervasive transcription and, therefore, having no apparent functional role, accumulating evidence indicates that such processes can have major repercussions on the activities of neighbouring genes [13]. Indeed, the effect of lncRNA transcription on gene activation or repression has been reported by a few studies [14,53–55]. Interestingly, this phenomenon does not seem to be restricted to lncRNAs but also extends to protein-coding mRNAs and, potentially, to all transcriptional events [20]. Gene-associated RNAs can recruit chromatin remodelers that are able to to maintain an open chromatin state or act as binding platforms for protein complexes at gene-proximal sites, such as the transcriptional factor Yin and Yang 1 (YY1) [56] and the MLL complex subunit WD repeat-containing 5 (WDR5) [57]. As a result, transcription upstream and downstream of annotated genes that we identified in this study might be functionally important for maintaining an open chromatin state and for the correct expression of neighbouring genes. That these transcripts are tightly associated with chromatin and are rapidly degraded by nuclear surveillance processes suggest their functions do not go beyond the course of transcription. For example, DoGs are highly sensitive to XRN2-mediated degradation; it has been previously reported that this protein promotes transcriptional termination at protein-coding genes via the torpedo mechanism model, in which the exonuclease degrades the gene-associated RNA until it reaches the elongation complex so causing its termination [58–60]. Hence, transcription of very long DoGs might underlie a longer engagement of RNA Pol II and, consequently, its inability to readily detach from DNA and restart transcription elsewhere and its contribution to maintain chromatin in an open state.

These mechanisms of transcription-associated chromatin regulation are not necessarily confined to intergenic regions linked to previously annotated genes but might be broadened to independent TUs. However, since these regions are usually found in gene-poor portions of the genome, their products are more likely to exert their functional role in *trans*. This and their similarities in terms of transcriptional activity, chromatin state and degradation patterns to lncRNAs support the hypothesis that independent TUs could be novel lncRNA loci.

## Conclusions

In summary, we assembled publicly available RNA-seq data to identify and classify intergenic transcripts based on their expression and location relative to annotated genes. We showed that gene-associated and independent RNAs have characteristic patterns of transcription and that they are highly sensitive to nuclear degradation processes. Our data are consistent with recently reported chromatin remodelling and gene expression regulatory mechanisms associated with transcription. Collectively, the results expand the current categories in gene annotation and provide the tools to further investigate the underappreciated role of intergenic transcription as a function of gene expression and regulation.

## Methods

### Reads alignment and post-processing

Sequencing quality checks were performed on all experiments using FastQC [61]. Adaptor sequences were removed using TrimGalore (v0.4.4_dev) [62] with default parameters. Reads were filtered against human rRNA and tRNA sequences obtained from the NCBI using Bowtie2 (v2.3.3.1) [63] with the option --sensitive-local. Reads that failed to align were mapped with STAR (v2.5.3a) [64] to UCSC hg38/GRCh38 genome assembly using GENCODE (v27) gene annotation [34] as reference, with the following parameters: *--twopassMode Basic --alignSJoverhangMin 8 --alignSJDBoverhangMin 1 --sjdbScore 1 --outFilterMultimapNmax 1 --outFilterMismatchNmax 999 --outFilterMismatchNoverReadLmax 0.04 --outFilterType BySJout --outSAMattributes All --outSAMtype BAM SortedByCoordinate*, and specific options for gapped (*--alignIntronMin 20 --alignIntronMax 1000000 --alignMatesGapMax 1000000*) and ungapped (*--alignIntronMax 1 --alignMatesGapMax 300*) alignments. PCR duplicates were removed using Picard MarkDuplicates (v2.18.3) with default parameters. Quantification of expression was performed using QoRTs (v1.3.0) [65] and the GENCODE (v27) gene annotation [34].

### Genomic coverage tracks

Deduplicated unique alignments were converted to stranded normalised coverage bigWig files using deeptools (v3.0.2) [66] with *--normalizeUsing CPM --binSize 20 --smoothLength 60* options, and *--filterRNAstrand* for the selection of forward and reverse strands.

### De novo transcriptome assembly

Deduplicated uniquely mapped reads were assembled into a *de novo* annotation GTF using StringTie (v1.3.4c) [33] with the GENCODE (v27) gene annotation [34] as reference, and the following parameters: *-f 0.2 -g 100 -j 3 -t*. The individual annotation GTFs from all wild-type RNA-seq datasets (no treatment condition was used to annotate the intergenic regions) were then used as input for StringTie with*--merge* option to generate a non-redundant set of predicted transcripts. The output, which consists of a GTF file with merged gene models, was filtered using the gffcompare utility [67] with *-C* option to discard predicted transcripts that were fully contained within larger annotated regions.

### Identification of intergenic transcriptional units

A custom R script was used to process the GTF file generated as described above. The script performs several steps, the first of which is the discrimination of the purely intergenic regions (*i.e.*, defined using the sequencing data) from the known features (*i.e.*, already present in the GENCODE gene annotation). This operation is performed by the *setdiff()* function from the *GenomicFeatures* R package [68] on the gffcompare-generated GTF and GENCODE reference annotation files. Intergenic regions with length ≤1 kb are discarded, while the remaining are further divided into “gene-associated” and “independent” transcriptional units (“gene-associated TUs” and “independent TUs”, respectively) based on whether they originate from annotated genes, thus showing transcriptional continuity with the gene body, or from regions devoid of annotated features, and therefore they are considered independent events of transcription. Gene-associated transcriptional units are assigned to different sub-groups depending on their position and connection to neighbouring gene(s):

- UpstreamOfGene (UoG): the unit is located upstream of the associated gene;
- DownstreamOfGene (DoG): the unit is located downstream of the associated gene;
- LinkerOfGenes (LoG): the unit is located between two genes, and transcriptionally associated with them.

To confirm the co-occurrence of the annotated gene(s) and gene-associated features, their expression is re-assessed across the RNA-seq datasets. Features are considered co-transcribed if expressed (TPM ≥ 1) in the same cell line and in at least two datasets. At this level, the categorisation is also re-evaluated and, if necessary, TUs can be re-assigned to the proper sub-group (*e.g.*, a LinkerOfGenes TU whose downstream gene is not expressed will become a DownstreamOfGene TU).

### Splicing analysis

Deduplicated unique alignments were parse using samtools [69] *view* and gapped alignments (*i.e.*, reads encompassing known or putative splice junctions) we extracted based on their CIGAR information (*i.e.*, whether or not it contained ‘N’). Reads were then assigned to ‘long ncRNA’ or ‘independent TU’ features using the *countOverlaps()* function from the *GenomicFeatures* R package [68]. For each dataset the fraction of junction reads was calculated over the total number of deduplicated unique reads.

### Selection of HeLa TUs and metadata profiles

Since the large majority of data available for validation derived from HeLa cells, we decided to focus our analysis of intergenic features only to those expressed in this cell line. Therefore, we generated a set of annotated genes and gene-associated and independent TUs where each feature had average expression ≥1 TPM across the HeLa RNA-seq datasets. In addition, we required the gene-associated features to be connected to annotated protein-coding genes, thus reducing the chance to include poorly annotated genes for which start and end genomic coordinates are not reliable (e.g., pseudogenes). We retained only the independent TUs located ≥10 kb from any annotated feature on the same strand orientation, to ensure that their transcription is not directly linked to known genes. Finally, we discarded features with length <5 kb to avoid signal overlaps between start and end positions in metadata profiles.

The metadata profiles were generated using the CPM normalised coverage bigWig files (see ‘*Genomic coverage tracks*’ section) and a custom wrapper of the *ScoreMatrixBin()* function from the *genomation* R package [70]. The wrapper function is used to facilitate strand splitting, centering and resizing (*i.e.*, ±5 kb from region start or end position), binning (*i.e.*, 200 bins over the 10kb window) and normalisation and averaging of the signal. When not specified in the figure legend, normalisation was performed by dividing the bins of each feature (or group of features in case of paired annotated gene and its gene-associated TU) by the value of the bin with the higher count across the region.

### Epigenetic modification profiles

We collected the ‘fold change over control’ and merged replicates ChIP-seq bigWig files from ENCODE. The list of epigenetic modifications and associated accession numbers can be found in Supplementary Table 1. The ChIP-seq signals across the regions of interest were calculated using the wrapper function described in the previous section.

### CAGE peaks profiles

We retrieved the hg38 CAGE reprocessed data [36] from the FANTOM Consortium [71]. The density of the CAGE peaks (phase 1 and 2) was calculated using the wrapper function described in the ‘*Selection of HeLa TUs and metadata profiles*’ section, without applying any normalisation.

### Quantification of expression and degradation

We collected the wild-type/untreated and several proteins knockdowns from different sources (see Supplementary Table 1). The datasets were processed as described in the ‘*Reads alignment and post-processing*’ section. Deduplicated uniquely mapped reads were loaded into R using the *GenomicAlignments* R package [68], and the expression of the features quantified with the *summarizeOverlaps* function. The *estimateSizeFactorForMatrix* function from the DESeq2 R package [72] was used to normalised the feature counts for each group of experiments. The ggpubr R package was used to visualise the results and perform the statistical tests (*i.e.*, two-sided Wilcoxon rank sum test).

## Declarations

### Ethics approval and consent to participate

Not applicable

### Consent for publication

Not applicable

### Availability of data and materials

The datasets analysed in the current study are available from public domain resources. The list of accession numbers is included in this published article and its supplementary information files.

### Authors’ contributions

F.A., J.U. and N.M.L. conceived the original idea and managed the project; F.A. developed bioinformatic pipeline; F.A. and J.U. planned the computational analyses; F.A. performed the computational analyses; J.Z., J.A., J.U. and N.M.L. provided valuable feedback on computational analyses and provided interpretation of the results; all the authors provided critical feedback and helped to shape research, analysis and manuscript; F.A. and J.Z. wrote the manuscript with valuable feedback from all authors.

## Acknowledgements

The authors are grateful to C. Pederiva and A. M. Chakrabarti for valuable assistance; and C. Pederiva for comments on this manuscript and for valuable advice.

## Authors’ information

F.A. is currently employed as a Postdoctoral Researcher at Science for Life Laboratory, Department of Medical Biochemistry and Biophysics, Karolinska Institutet, Stockholm, Sweden.

---

**Supplementary Figure 1. Summary of characteristics of gene-associated and independent TUs.** A) Distribution of genomic distances between independent TUs and the nearest annotated gene. B) Distribution of proportions of spliced reads among annotated lncRNAs and independent TUs; each data point represents an RNA-seq dataset included in this study. C) Gene-associated TUs nearest annotated gene type; D) Gene-associated and intergenic TUs expression against their nearest annotated gene expression.

**Supplementary Figure 2. Meta-profiles of transcriptional measurements around LoGs.** Meta-profiles of transcriptional measurements plotted relative to the start and end positions of LoGs and their associated protein-coding genes start positions. Panels A-D as in Figure 3.

**Supplementary Figure 3. NET-seq meta-profiles for different Pol II CTD modifications around TUs.** RNA polymerase II and its CTD modifications (NET-seq) occupancy profiles across: A) UoG regions and connected protein-coding genes start positions (left) and protein-coding genes and connected DoG regions end positions (right); B) upstream gene and LoG start (left) and LoG and downstream gene end (right) positions; C) long non-coding RNA genes and independent transcriptional units start (left) and end (right) positions.

**Supplementary Figure 4. Enhancer epigenetic signature profiles.** Enhancer-associated histone marks profiles across: A) UoG regions and connected protein-coding genes start positions (left) and protein-coding genes and connected DoG regions end positions (right); B) upstream gene and LoG start (left) and LoG and downstream gene end (right) positions; C) long non-coding RNA genes and independent transcriptional units start (left) and end (right) positions.

**Supplementary Figure 5. Impact of nuclease-depletion on gene-associated TU expression.** Panels A-D as in Figure 5.

**Supplementary Table 1. Datasets used in this study.** Annotation (sheet 1), validation (sheet 2) and histone modifications samples, including accession numbers and mapping metrics.

