## Supplementary material for "Intergenic RNA mainly derives from nascent transcripts of known genes"

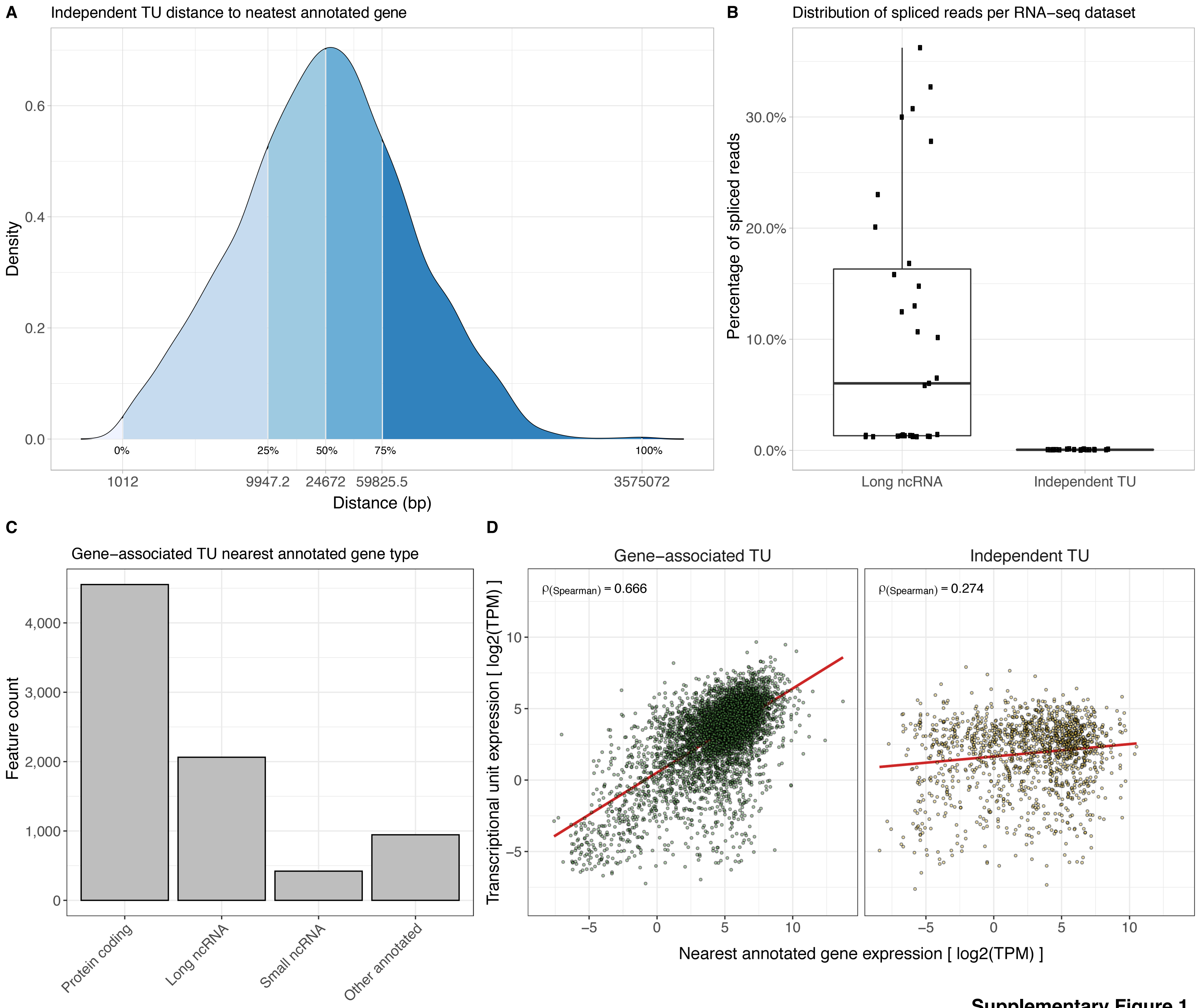


**Supplementary Figure 1**

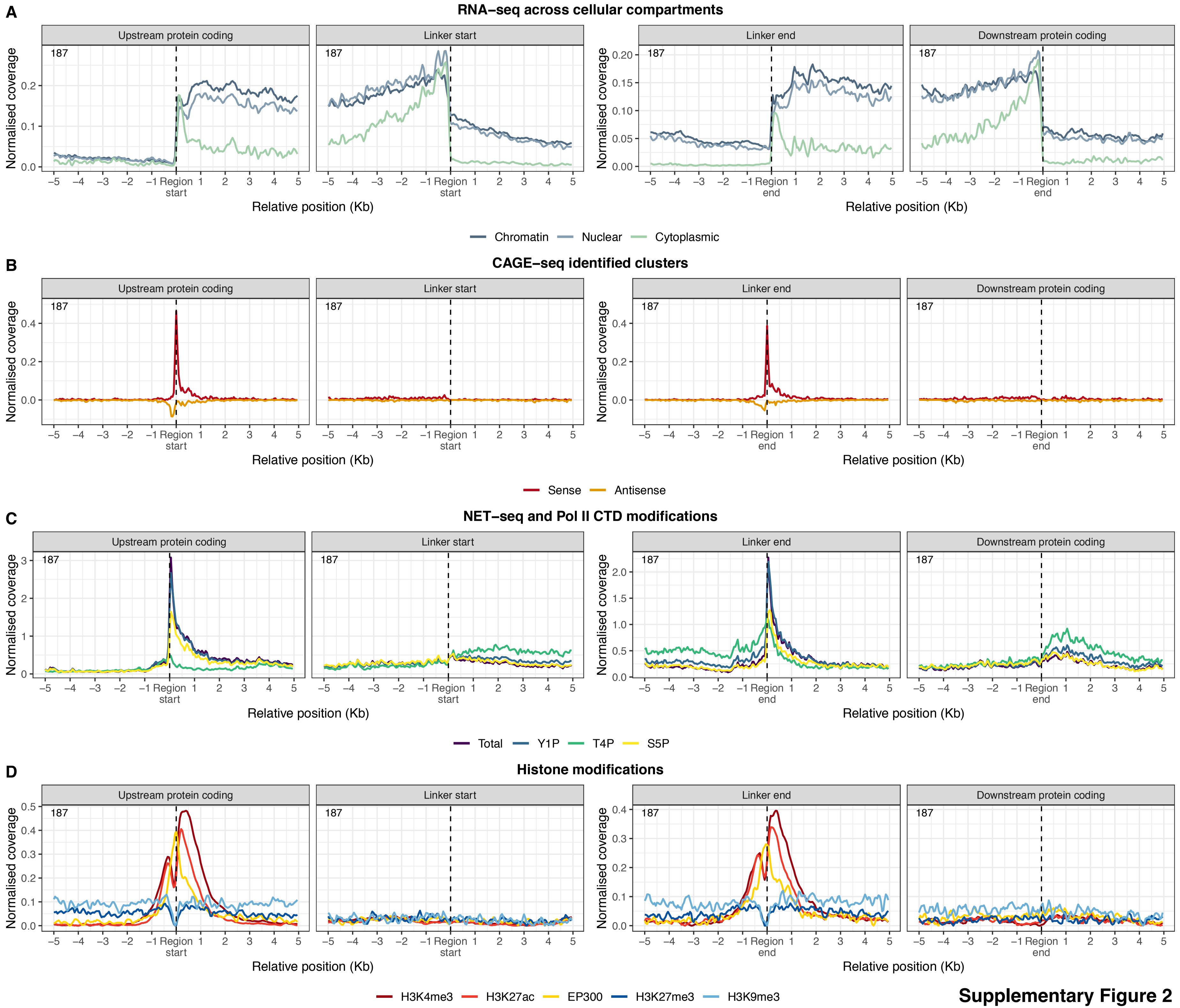


Supplementary Figure 2

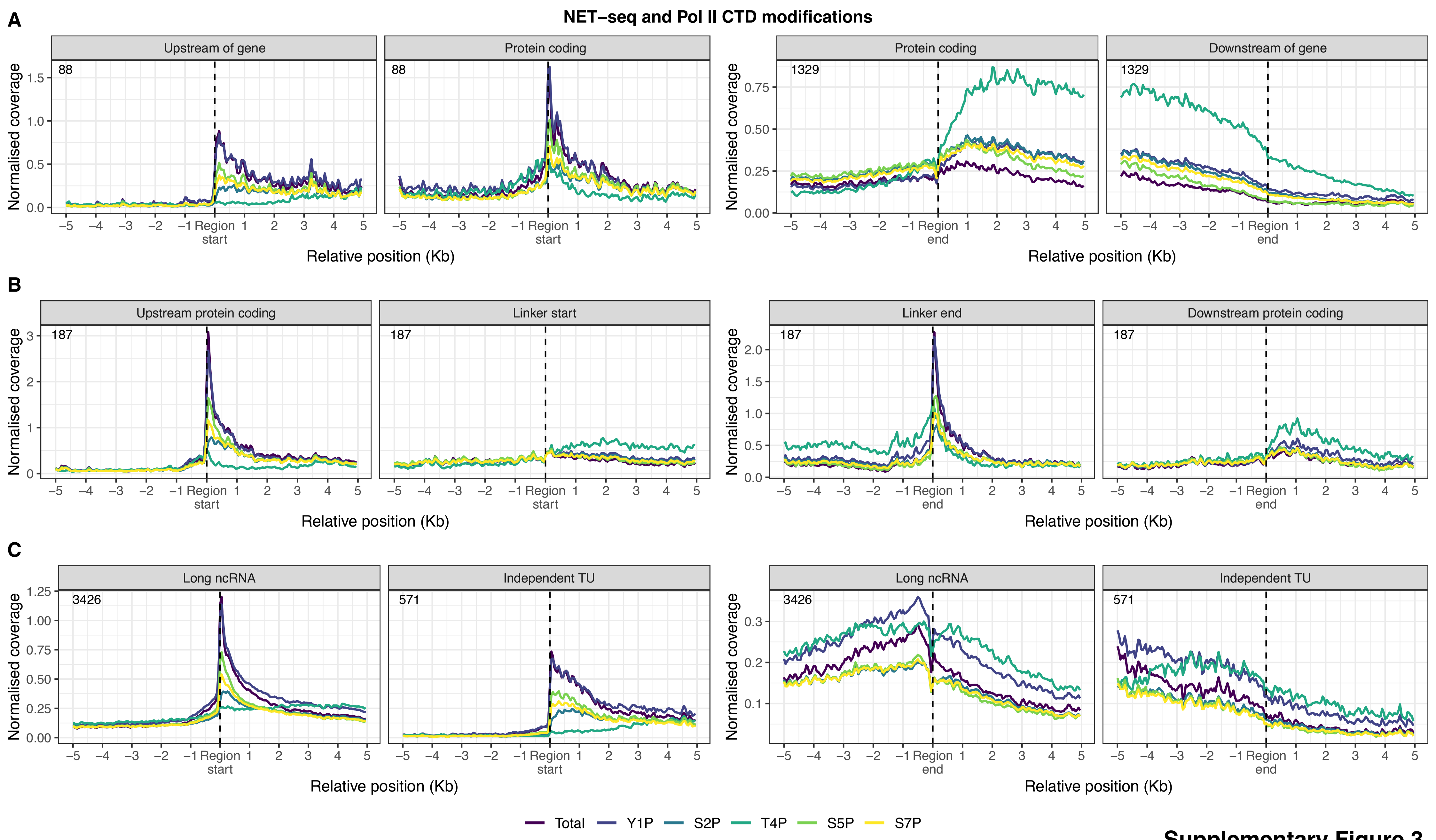


Supplementary Figure 3

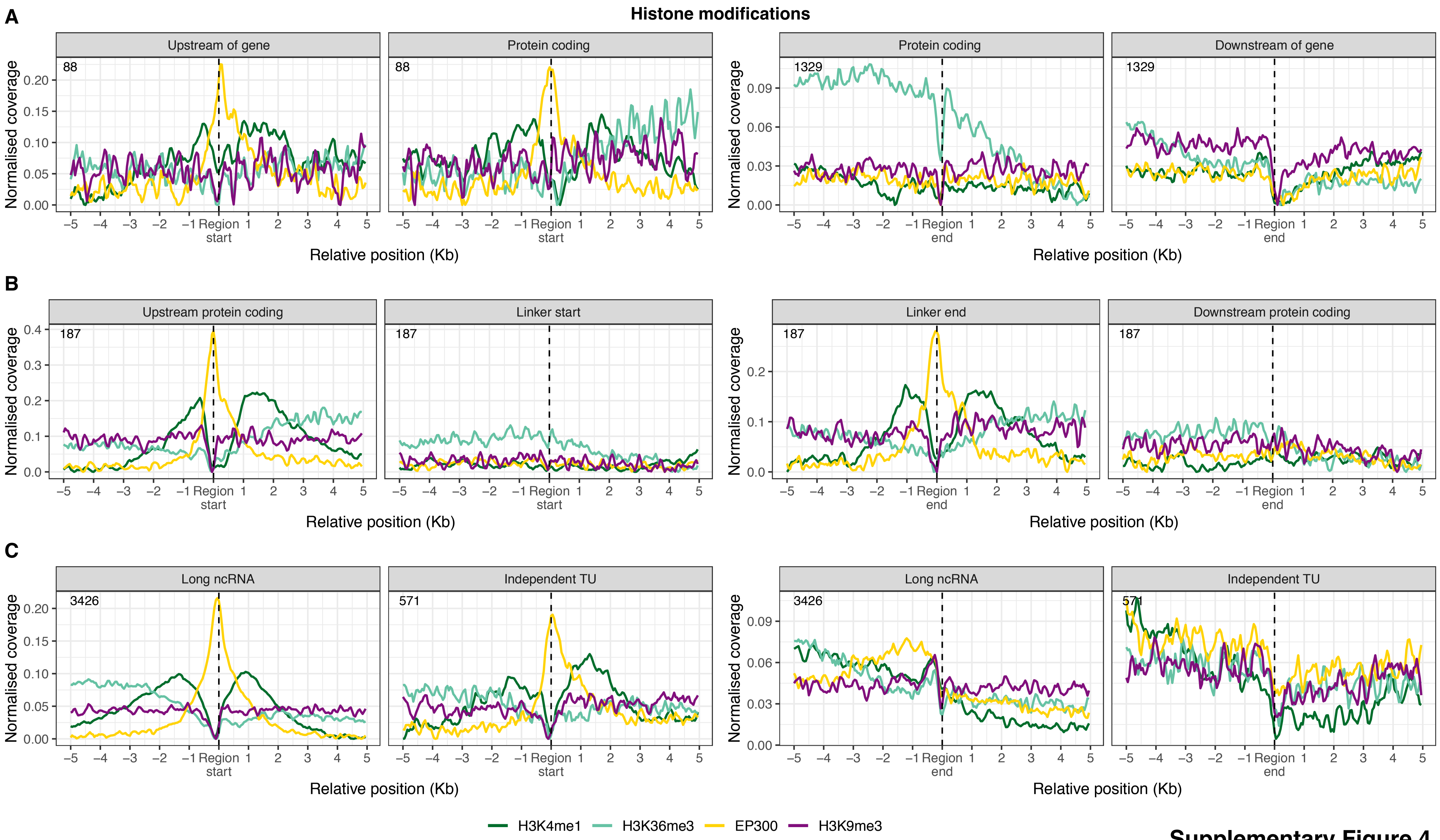


**Supplementary Figure 4**

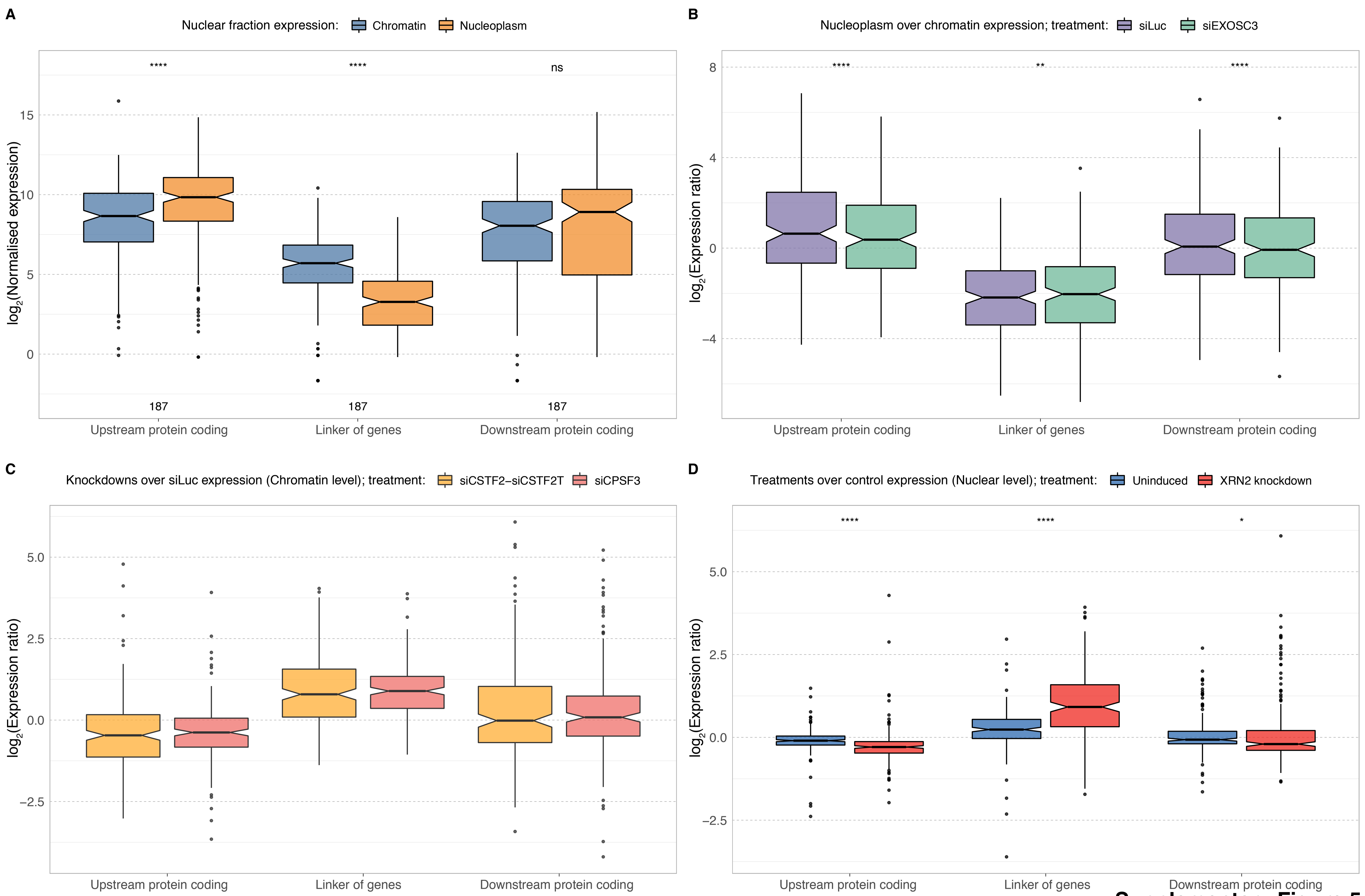


**Supplementary Figure 5**
